# Transgenic Zebrafish Models Reveal Distinct Molecular Mechanisms for Cataract-linked αA-Crystallin Mutants

**DOI:** 10.1101/364125

**Authors:** Shu-Yu Wu, Ping Zou, Sanjay Mishra, Hassane S Mchaourab

## Abstract

Mutations in the small heat shock proteins α-crystallins have been linked to autosomal dominant cataracts in humans. Extensive studies *in vitro* have revealed a spectrum of alterations to the structure and function of these proteins including shifts in the size of the oligomer, modulation of subunit exchange and modification of their affinity to client proteins. Although mouse models of these mutants were instrumental in identifying changes in cellular proliferation and lens development, a direct comparative analysis of their effects on lens proteostasis has not been performed. Here, we have transgenically expressed cataract-linked mutants of αA- and αB-crystallin in the zebrafish lens to dissect the underlying molecular changes that contribute to the loss of lens optical properties. Zebrafish lines expressing these mutants displayed a range of morphological lens defects. Phenotype penetrance and severity were dependent on the mutation even in fish lines lacking endogenous α-crystallin. The mechanistic origins of these differences were investigated by the transgenic co-expression of a destabilized human *γ*D-crystallin mutant. We found that the R49C but not the R116C mutant of αA-crystallin promoted aggregation of γD-crystallin, although both mutants have similar affinity to client proteins *in vitro*. Our working model attributes these differences to the propensity of R49C, located in the buried N-terminal domain of αA-crystallin, to disulfide crosslinking as previously demonstrated *in vitro*. Our findings complement and extend previous work in mouse models and emphasize the need of investigating chaperone/client protein interactions in appropriate cellular context.

## Introduction

In its most common form, age-related cataract is an opacity of the lens characterized by the formation of protein aggregates that scatter light [1]. Protein aggregation is driven by the progressive insolubilization of lens proteins as a consequence of age-dependent changes to their sequences and structures [2, 3]. Terminally differentiated lens fiber cells, devoid of cellular machineries that repair and turn proteins over, rely on two small heat shock proteins (sHSPs), αA-and αB-crystallin, which function as chaperones by sequestering thermodynamically destabilized client proteins to inhibit aggregation [2–5]. The prevailing model of age-related cataract posits that the chaperone capacity of α-crystallins is titrated out by binding of damaged lens proteins as well as truncation and insolubilization of the sHSPs themselves [6–8]. Detailed models of stable and transient interactions with model client proteins revealed activation mechanisms of α-crystallins and defined the energetics of the chaperone interactions [9–17].

Consistent with a central role in lens development and transparency, point mutants of α-crystallins have been associated with hereditary cataracts [5, 18]. Multiple studies dissected how the mutations affect the interactions with client proteins *in vitro* [19–22]. Koteiche and Mchaourab demonstrated that two αA-crystallin substitutions, R49C and R116C, lead to a 100-fold increase in the affinity to the client protein T4 lysozyme (T4L) [11]. Interpreted in the framework of a model where the affinity of the chaperone is thermodynamically coupled to the folding equilibrium of T4L, these findings supported the conclusion that the mutants can behave as “unfoldases” [23]. The mutated chaperones are predicted to bind undamaged proteins, which promotes their unfolding, and consequently leads to saturation of the binding sites and the formation of large chaperone/protein aggregates [23]. A similar conclusion was reached for the αB-crystallin R120G, a human mutation which has been associated with cardiomyopathy [20].

Mouse models of αA- and αB-crystallin mutants associated with hereditary cataract delineated a number of important cellular changes and shed light on previously unappreciated physiological roles for these proteins in apoptosis and genomic maintenance [24–27]. However, due to inherent limitations of the mouse model, a direct analysis of how modification of chaperone/client protein interactions affects lens proteostasis is not straightforward. Studies of the knock-in model of αA-R49C suggested changes in these interactions as well as in the level of insolubilization of lens proteins [26]. However, the former was only observed in the heterozygotes, which paradoxically had a moderate cataract phenotype. Thus, while these preliminary observations are consistent with the model of Koteiche and Mchaourab [11], a more detailed interrogation of these interactions in the lens is needed.

To establish a mechanistic link between *in vitro* models of α-crystallin chaperone function and their roles in lens aging, we have utilized zebrafish (*Danio rerio*) as an experimental model organism. A number of advantages makes zebrafish ideal for such an endeavor including ease and cost of rearing the animals and the size of the embryo clutches [28, 29]. Moreover, the transparency of the embryos during the first few days of their development and their relatively large eyes, which are functional at 3 days post fertilization (dpf), enable the examination of lens gross morphology by bright field microscopy [30–34]. The structure and function of the zebrafish adult lens parallel those of the mature human lens with respect to symmetry, refraction, transparency and optical function [29, 35, 36]. Previous work from our laboratory demonstrated that both α-crystallins are required for lens development [32, 37]. Furthermore, we established the utility of the zebrafish as a model for lens proteostasis by demonstrating a correlation between the level of gross lens defects and the expression of progressively destabilized mutants of human *γ*D-crystallin [33].

Here, we transgenically expressed two different mutants of αA-crystallins associated with hereditary cataract and the cardiomyopathy-linked R120G mutant of αB-crystallin in the zebrafish lens. In addition to characterizing the phenotypes of the transgenic lines, we challenged the proteostasis network with the concurrent expression of a destabilized mutant of *γ*D-crystallin. Initially identified in γB-crystallin from a cataract mouse model [38], the I4F substitution has been studied extensively *in vitro* [39, 40]. The relatively minor destabilization and aggregation propensity of the mutant makes it ideal to investigate chaperone interactions with α-crystallin. Together, the results presented here reveal differences in the penetrance and severity of the phenotypes caused by the transgenic expression of the α-crystallin mutants. We demonstrate that the more deleterious effects of R49C are the result of its promoting the aggregation of lens proteins as predicted from the thermodynamic coupling model [11, 41].

## Materials and Methods

### Zebrafish maintenance and breeding

AB wild-type strain zebrafish (*Danio rerio*) were used. The embryos were obtained by natural spawning and raised at 28.5°C on a 14/10 hour light/dark cycle in 0.3x Danieau water containing 0.003% PTU (w/v) to prevent pigment formation. Embryos were staged according to their ages (dpf; days post-fertilization). The following mutant and transgenic fish lines were used: *cryaa*^vu532^ [32], *cryaba*^vu612^; *cryabb*^vu613^[37], Tg(*cryaa*:*Rno.Cryaa*_*R49C*,*myl7*:Cerulean)^vu615Tg^, Tg(*cryaa*:*Rno.Cryaa*_*R116C*,*myl7*:Cerulean)^vu616Tg^, Tg(*cryaa*:*Mmu.Cryab*_*R120G*,*myl7*:Cerulean)^vu617Tg^ (this study), Tg(*cryaa*:Gal4vp16)^mw46Tg^; Tg(*UAS*:GFP)^kca33Tg^ [42]. All animal procedures were verified and approved by the Vanderbilt University Institutional Animal Care and Use Committee.

### Zebrafish transgenesis

To establish the transgenic zebrafish expressing rat (*Rno*) *Cryaa* gene (*Rno.Cryaa*) specifically in the lens, Tg(*cryaa:Rno.Cryaa*,*myl7:Cerulean*) was constructed by inserting mutated *Rno.Cryaa* cDNA (by site-mutagenesis; R49C and R116C) downstream of zebrafish *cryaa* promoter (1.2 kb)[43] in the pT2HBLR vector that was also contains *myl7* promoter-driven Cerulean as the selection marker (i.e. in the heart). Tol2 mediated transgenesis were performed as previously descried [32] for the *Rno.Cryaa* (R49C and R116C) transgenes. The same protocol was followed for generating the mouse *cryab* R120G variant (*Mmu.Cryab_R120G*) transgene. At least two founder lines (F0) for each construct were screened and out-crossed to established stable F1 generations. Each F1 line was propagated and raised into F2 and F3 generations. The lens defects data collected were from F3 or F4 embryos, and similar penetrance of lens defects was observed in individual stable lines that express the same α-crystallin constructs.

### Cell death assays

Embryos were fixed overnight at 4°C in 4% paraformaldehyde in PBS, dehydrated with 100% MeOH and stored in −20°C. The procedures of TUNEL staining were carried out following the manufacturer’s suggested protocol (In Situ Cell Death Detection Kit, TMR red; Sigma-Aldrich #12156792910).

### Microscopy and image processing

Lenses of live embryos in 0.3x Daneau water with PTU/tricane were analyzed by bright field microscopy (Zeiss Axiovert 200) at 4dpf and graded into three classes depending on the severity of lens defects as defined in our previous study [32]. Fluorescence images were taken with Zeiss AxioZoom.V16 microscope. Differential interference contrast (DIC) and reflectance analysis were performed by Zeiss LSM510 inverted confocal microscope with λem=488 on 4dpf embryos which were embedded in 2.5% methylcellulose. DIC images were taken with a 20x objective lens and used as a general locator for reflectance analysis. Reflected light was analyzed by digital image analysis program Image J.

### Statistics

Differences among groups were analyzed by Student’s *t*-test. Data are shown as means ± standard error (SE). Statistical significance was accepted when p < 0.05.

## Results

### Transgenic expression of αΑ-crystallin variants induce embryonic lens defects in zebrafish

To investigate the *in vivo* interactions between cataract-linked αΑ-crystallin mutants and lens chaperones in the context of the native environment of the lens fiber cells, we generated transgenic zebrafish lines that express the αΑ-crystallin mutants R49C and R116C (*Tg[cryaa:Rno.Cryaa_R49C] and Tg[cryaa:Rno.Cryaa_R116C]*; hereafter referred to as“*αΑ-R49C*” and “*αΑ-116C*” for simplicity). Using zebrafish transgenesis protocols established previously [32], the two αΑ-crystallin variants were specifically expressed in the lens under the control of the zebrafish *cryaa* promoter with *myl7* promoter-driven Cerulean fluorescent protein in the heart as a convenient selection marker for transgenic animals.

Expression of both αΑ-crystallin variants led to various degrees of embryonic lens abnormalities that were readily visible starting at 3dpf (Figure 1B), without affecting the overall morphology of the embryos (Figure 1A). The nature of these defects was similar to those previously described for α-crystallin deficient [32, 37] or γD-crystallin mutant transgenic lines [33]. Defective lenses exhibited phenotypic features that were either spherical, shiny crystal-like droplets spread sporadically across the lens (minor defects), or frequent droplets covering a large fraction of the lens as well as occasional large irregular protuberances located in the center of the lens (major defects), all of which could lead to opacity and changes in light scattering. For each αΑ-crystallin mutant, we screened a large number of embryos and scored their lens phenotype based on the severity (Figure 1B). While the expression of both mutants led to a considerable fraction of embryos displaying lens defects, the data suggest a more deleterious consequence for the R49C mutation both in the penetrance and severity of the phenotype.

**Figure 1.**
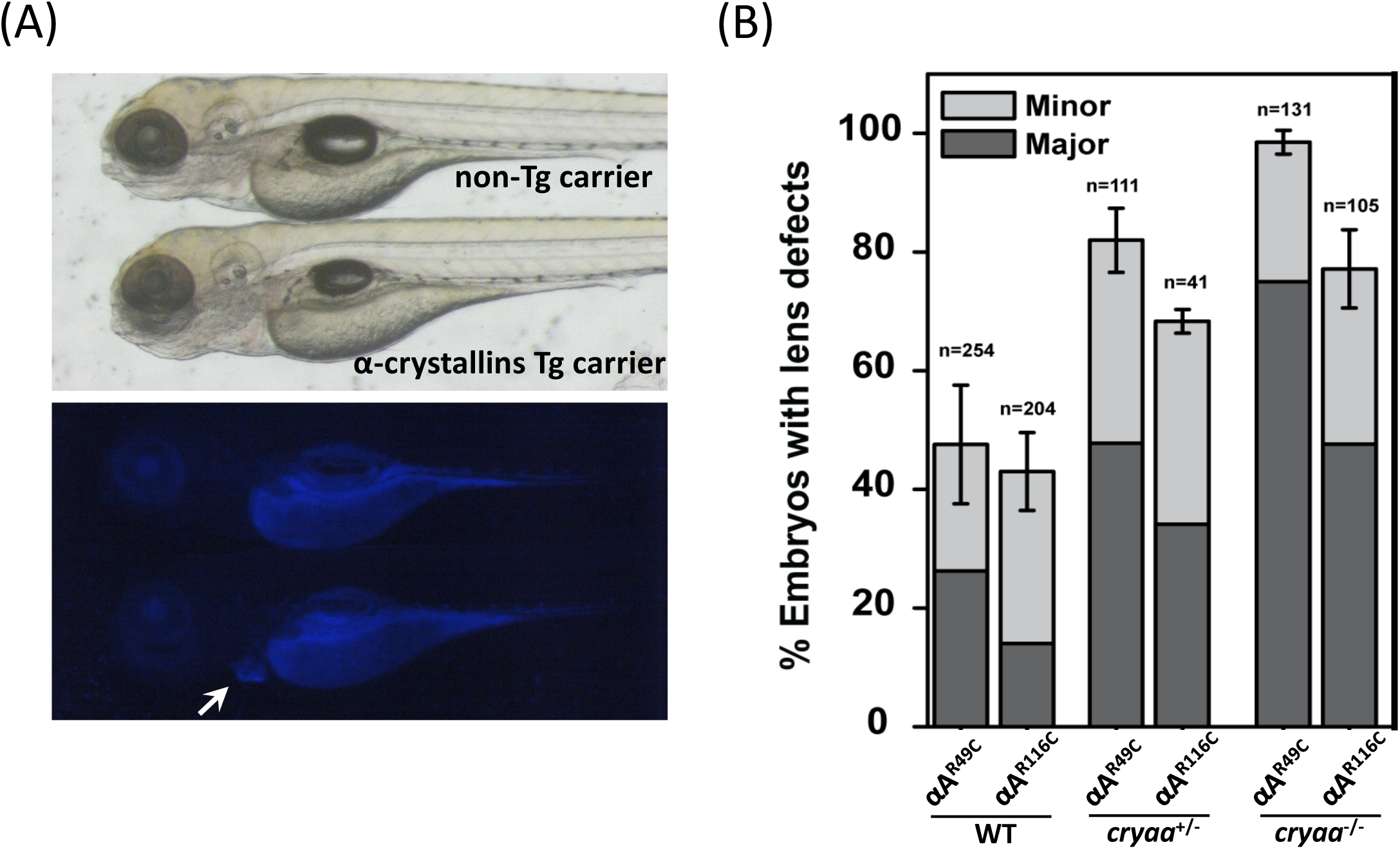
Transgenic expression of murine *Cryaa* mutants induced lens defects in 4dpf zebrafish embryos. (A) The lens-specific murine *Cryaa* transgenes displayed *Cerulean* marker in the heart. *Upper Panel*, DIC image. *Lower panel*, a *white arrow* marks the *Cerulean* marker in the heart. No overall morphology changes were observed in all transgenic carriers. (B) Embryos of lens-specific *Rno.Cryaa* transgenes (R49C and R116C mutations) displayed lens defects with various penetrance. Removal of endogenous *cryaa* alleles exacerbated the lens defects.

### Genetic dosage of αΑ-crystallin determines the level of lens defects induced by the mutants

The R49C and R116C mutations of human αΑ-crystallin cause congenital cataracts in an autosomal dominant fashion, while only 40~50 % of the embryos show lens defects in the transgenic fish lines expressing these mutant forms of αΑ-crystallin. The gene dosage of αΑ-crystallin has been shown to impact the penetrance and/or severity of lens phenotypes, as demonstrated earlier in *cryaa* zebrafish mutants [32] as well as the *Cryaa*-R49C knock-in mouse model [26], where the lenses from *Cryaa*-R49C homozygotes (equivalent of losing two copies of wild-type *Cryaa*) showed more severe lens opacity.

To investigate if the relatively moderate penetrance of lens defects was due to the expression of the endogenous zebrafish *cryaa* gene (i.e. wild-type) in these transgenic lines, we generated heterozygote αΑ-crystallin mutants. For this purpose, the *αΑ-R49C* and *αΑ-R116C* transgenic lines were crossed into *cryaa* null mutants [32] to create genetic conditions more closely mimicking the human autosomal dominant CRYAA mutations. The loss of one endogenous *cryaa* allele (i.e. *cryaa*^+/−^), led to an increase in the percentage of lens defects from both αΑ-crystallin mutant transgenic lines, albeit not to 100% (Figure 1B). Loss of the other endogenous *cryaa* allele by backcrossing to generate αΑ-crystallin mutant transgenic lines under the *cryaa* null background (*cryaa*^−/−^) further increased the percentage of embryos with lens defects approximating 100% penetrance for *αΑ-R49C* and near 80% for *αΑ-R116C*.

Because cell death has been implicated in cataract phenotypes, including radiation-induced cataracts, diabetic retinopathy-associated cataracts [44–46] as well as age-related cataract [47], we tested for apoptotic cell death in the transgenic lines. TUNEL assay (Figure 2) did not show evidence of elevated apoptosis in the embryonic lens carrying αΑ-crystallin mutant transgenes when compared to WT. Therefore, cell death by apoptosis does not seem to play a major role in the embryonic lens defects in the αΑ-crystallin zebrafish transgenes, a result which differs the mouse αΑ-crystallin mutation knock-in and transgenic models [24, 26] where increase of cell death were observed in the postnatal lens.

**Figure 2.**
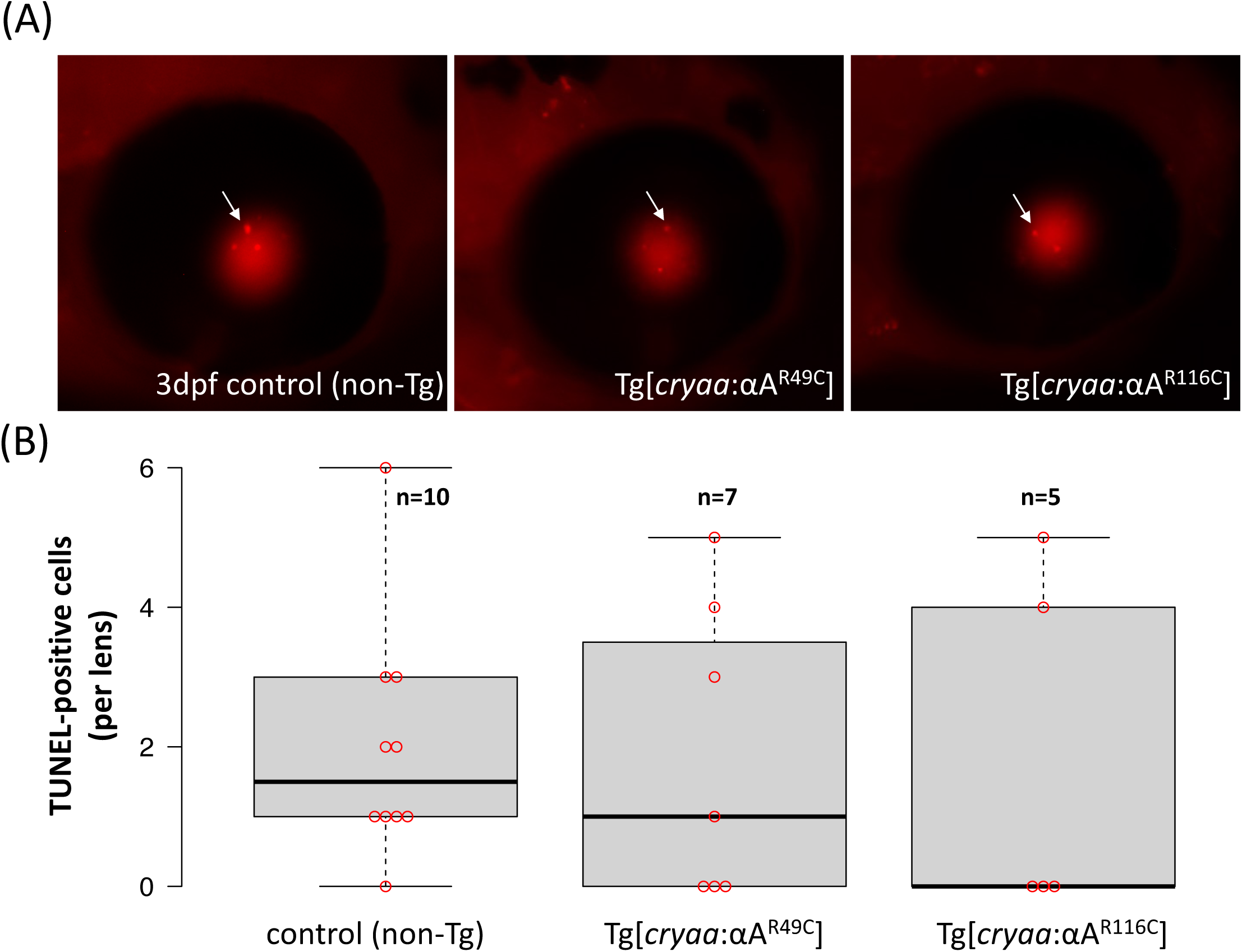
Apoptotic cell death was not induced by transgenic expression of murine *Cryaa* mutants. By the TUNEL staining (A), embryos of *Tg*(*cryaa:Rno.Cryaa_R49C*) *and Tg*(*cryaa:Rno.Cryaa_R116C*) did not show increase of apoptosis in the lens when compared to non-transgenic (WT) siblings (B).

### αA-R49C but not R116C increases the phenotype penetrance of a destabilized γ-crystallin mutant

The model of Koteiche and Mchaourab predicts that R49C and R116C would act to bind native lens proteins driving unfolding and aggregation [11, 48] and leading to formation of large particulates that scatter light. To test this model, we co-expressed each of the mutants with the human γD-crystallin variant I4F [33]. The I4F substitution reduces the free energy of unfolding reflecting a lower stability of the N-terminal domain [39]. In addition, the mutant exhibits a slow aggregation propensity [40]. Transgenic expression of γD-crystallin I4F mutant by itself resulted in moderate frequency (~30%) of major lens defects (*γD-I4F*; Figure 3), consistent with our previous study [33].

**Figure 3.**
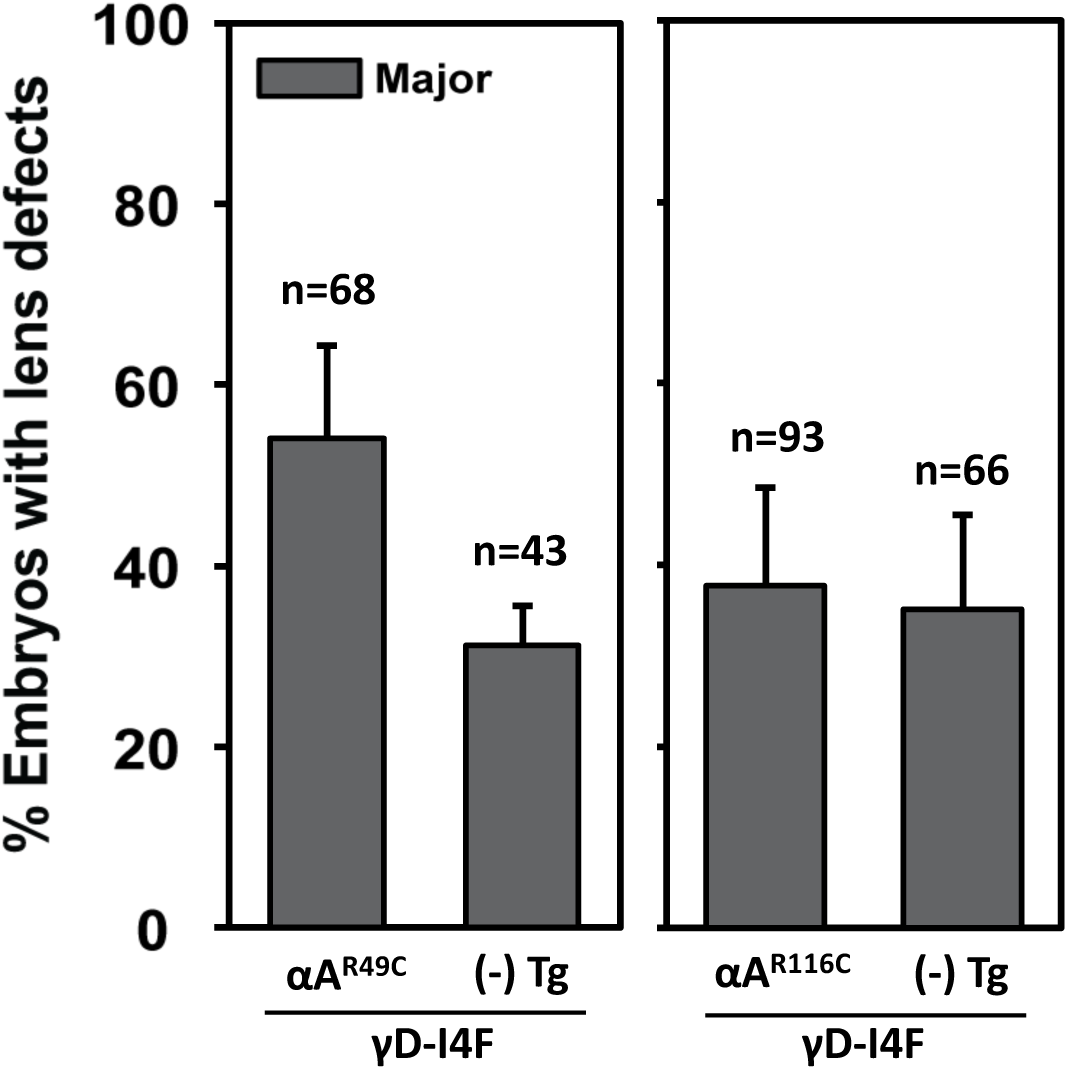
Synergistic effects on embryonic lens defects were selectively observed in transgenic co-expression of the human *CRYGD* mutant and murine *Cryaa* mutants. Compared to the siblings carrying single transgene, *Tg*(*cryaa:Hsa.CRYGD_I4F*), double transgenic embryos at 4dpf exhibited enhanced penetrance of lens defects when *Tg*(*cryaa:Rno.Cryaa_R49C*) was co-expressed, but not *Tg*(*cryaa:Rno.Cryaa_R116C*).

Co-expression of the *αΑ-R49C* transgene with *γD-I4F* (i.e. *αΑ-R49C*; *γD-I4F*) led to a synergetic effect significantly (p<0.05) increasing the percentage of major lens defects. In contrast, co-expression of *αΑ-R116C* did not significantly change the severity or penetrance of the lens defects (*αΑ-R116C*; *γD-I4F*), which remained comparable with single transgenic embryos (*γD-I4F*) (Figure 3).

### Expression of αA-R49C increases the propensity of destabilized γD-crystallin aggregation

In a previous study, we demonstrated that destabilized γD-crystallin mutant proteins formed aggregates in the zebrafish lens with a frequency that strongly correlated with the penetrance of lens defects [33]. Thus, the enhanced lens defects observed in the double transgenic embryos (Figure 3; *αΑ-R49C*; *γD-I4F*) could result from increased aggregation frequency of γD-I4F when the *αΑ-R49C* transgene was co-expressed.

We directly tested this model utilizing a similar experimental approach with the Gal4/UAS expression system to facilitate mosaic analysis (Figure 4A)[33]. By comparing the frequency of fluorescent punctate formation (i.e. mCherry-tagged γD-I4F) between transgenic carriers and non-transgenic embryos, we observed a significant difference between *αΑ-R49C* and *αΑ-R116C* transgenes in promoting γD-crystallin aggregation (Figure 4B). In the presence of the *αΑ-R49C* transgene, the frequency of γD-crystallin aggregation was higher by about 15~20% (Left panel); while no apparent difference was observed regardless the expression of *αΑ-R116C* (Right panel). As expected, the phenotype penetrance of *αΑ-R49C* is increased by the loss of endogenous αA-crystallin.

**Figure 4.**
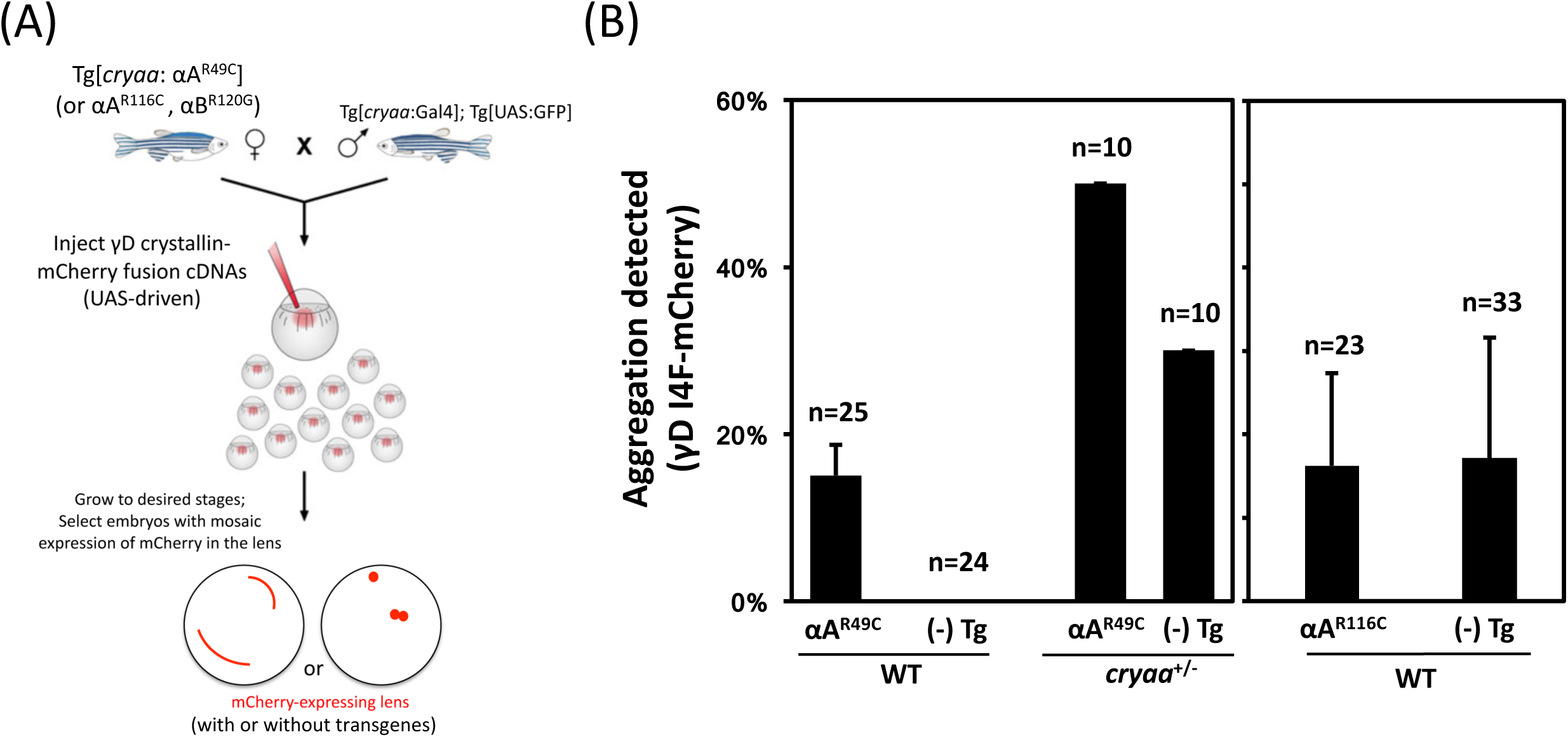
Differential modulation of γD-crystallin protein aggregation by transgenes of murine *Cryaa* mutants. (A) Schematic of the experiment utilizing the Gal4/UAS targeted gene expression system. Double transgenic line, Tg[*cryaa*:Gal4]; Tg[*USA*:GFP], were outcrossed to either *αA-R49C, cryaa^−/−^;αA-R49C, αA-R116C* (shown in Figure 4B) or *αB-R120G* (shown in Figure 5B) transgenic lines. Fertilized zygotes were injected with tol2 mRNA and UAS responder Tol2 constructs expressing fluorescently tagged human γD-crystallin (*Hsa.CRYGD_I4F-mCherry*). Embryos possessing lens fibers positive for mCherry mosaic expression were selected and imaged at 4dpf. (B) The frequency of embryos showing γD-crystallin_I4F-mCherry punctuates in the lens were significantly increased when *Tg*(*cryaa:Rno.Cryaa_R49C*) was co-expressed, but not with *Tg*(*cryaa:Rno.Cryaa_R116C*).

A higher aggregation frequency of γD-crystallin was observed for the non-transgenic embryos from the *αΑ-R116C* crosses compared to the ones from the *αΑ-R49C* crosses (Figure 4B). A number of potential reasons could contribute to this difference. First, the stochastic nature of mosaic analysis where the mCherry-tagged γD-crystallin cDNA constructs were randomly distributed in the lens fiber cells by microinjection probably led to different expression/distribution of γD-crystallin, even though the same nominal amount of DNA was injected into each 1-cell zygote. Consistent with this conjecture, a wide-range of positive F0 carriers was reported in the context of zebrafish transgenesis [49]. Second and of critical importance is the underappreciated phenomenon of cross-generational effects of genetic modifiers [50], which was also suggested to account for variations in lens defects severity in the progeny of *cryaa*^+/−^ adult lines in a previous study [32]. The adult transgenic *αΑ-R116C* and *αΑ-R49C* fish lines used here were neither phenotypically screened/selected before raised to adulthood, nor restricted for any particular sex (i.e. random usage of male and female fishes). Thus, the zygotes from *αΑ-R116C* and *αΑ-R49C* crosses may possess intrinsically different proteostasis capacity inherited from their parents, which would manifest as a difference in the basal level of protein aggregation.

### Expression of murine αB-crystallin R120G mutant induces zebrafish lens defects but does not promote γD-crystallin aggregation

αB-crystallin is a molecular chaperone involved in the cell response to stress in various tissues (see reviews in [51, 52]). Loss-of-function studies demonstrated that αB-crystallin is critical in maintaining lens proteostasis [37], and several missense mutations in human αB-crystallin gene are linked to hereditary cataracts, including P20S [53] and D140N [54]. The most-studied mutation, R120G, resides within the α-crystallin domain and is associated with desmin-related myopathy [20, 55].

Because αB-crystallin R120 is homologous to αA-crystallin R116 and the two mutations (R120G and R116C) have similar properties *in vitro*, we generated a transgenic line expressing the αB-crystallin R120G mutant specifically in the lens (*αB-R120G*). As expected *αB-R120G* induced lens defects with a moderate penetrance but the severity of lens phenotypes was mostly of the minor class (Figure 5A). Reducing endogenous αB-crystallin levels by crossing *αB-R120G* into zebrafish αB-crystallin null mutants (*cryaba*^−/−^; *cryabb*^−/−^) [37] led to significant enhancement of lens defects in both penetrance and severity (Figure 5A).

**Figure 5.**
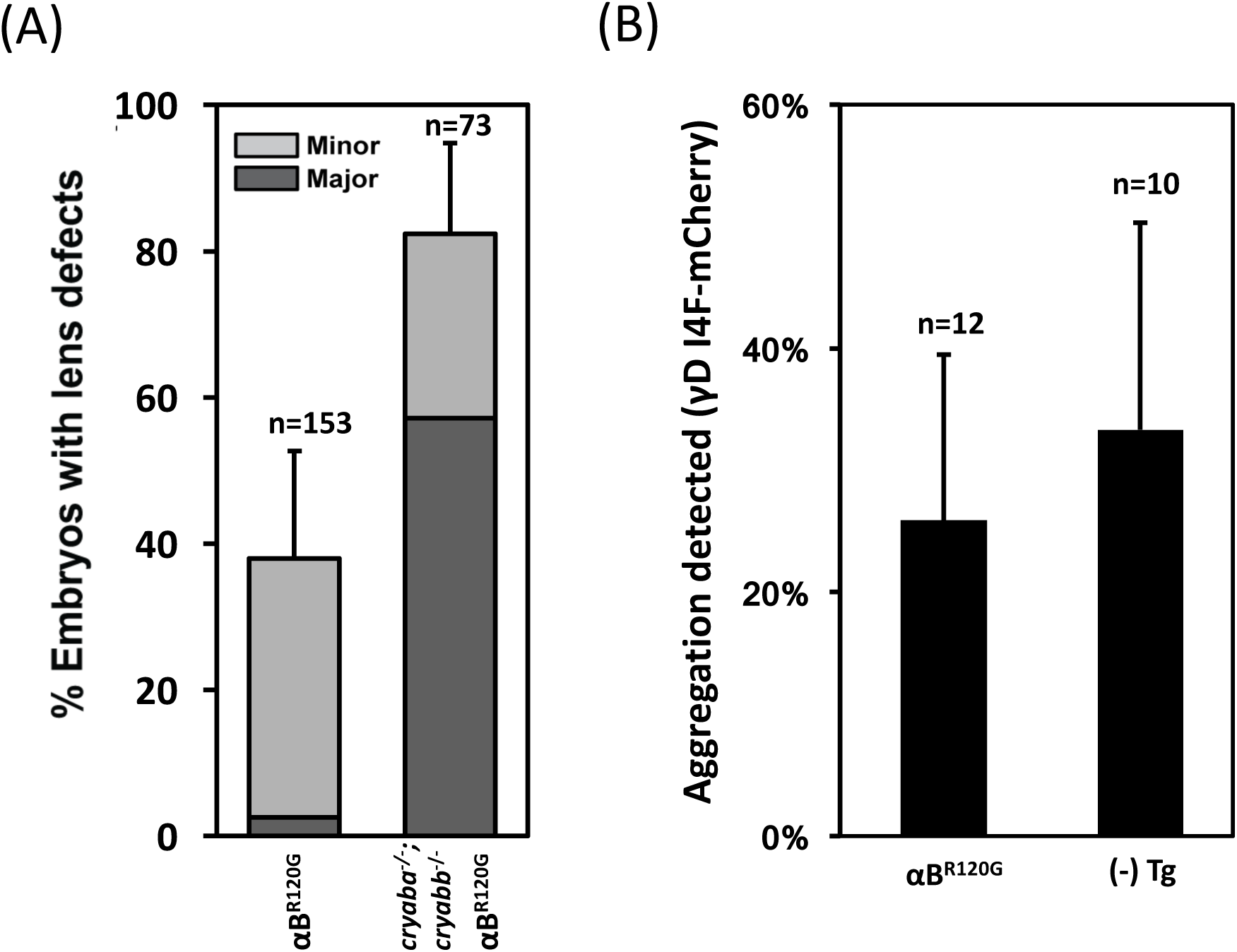
The effects on gD-crystallin protein aggregation by transgenes of murine *αB-crystallin R120G* mutant. (A) Transgenic expression of mouse *Cryab R120G* mutant induced lens defects in 4dpf zebrafish embryos, and lens defects were significantly exacerbated in the background of αΒ-crystallin null (*cryaba^−/−^; cryabb^−/−^*). (B) The frequency of embryos showing mCherry-tagged gD-I4F punctuates in the lens were not changed when co-expressed with *Tg*(*cryaa:Mmu.Cryab_R120G*).

If the nature of the substitution-i.e. glycine versus cysteine is not critical for the mechanism, the results of *αΑ-R116C* on the modulation of γD-crystallin aggregation predicts that *αB-R120G* would have a minimal role on the propensity of γD-crystallin aggregation. Indeed, transgenic expression of *αB-R120G* did not significantly increase the γD-I4F aggregation frequency when compared to non-transgenic siblings (Figure 5B). Together with the results from the αΑ-crystallin transgenic lines, this finding suggests that the tendency of R49C to increase γD-crystallin aggregation is intimately related to its position in the primary sequence.

## Discussion

The work reported here takes advantage of the power of zebrafish as a model organism to carry out the first direct comparison between cataract-linked mutants of αA-crystallin in a similar genetic background. R49C and R116C are two human autosomal-dominant mutations that have been associated with a wide range of effects on the structure and function of αA-crystallin. Similar to mouse models, expression of these mutants in the zebrafish lens leads to lens defects that have been previously demonstrated to cause increased light scattering [32, 33]. While loss of one copy of the WT protein (i.e. endogenous allele) accentuates the penetrance and severity of the phenotype, the R49C mutation was consistently more deleterious than the R116C mutation. We linked the differential effects of the two mutants to a higher propensity of R49C to induce protein aggregation. The two mutants were predicted to display similar chaperone properties from *in vitro* binding studies to the model client protein T4L [11]. Thus, our findings highlight the importance of *in vivo* investigation in validating mechanistic models.

How does co-expression of αA-R49C increase the propensity of γD-I4F to aggregate? Our thermodynamic coupling model predicts that an increase in the apparent affinity of the chaperone to the client protein would drive unfolding of latter by simple mass action. Because in the double transgenic line, the interaction with the γD-crystallin mutant is exclusive to the R49C mutant and the level of expression of the two proteins is expected to be largely similar, given that both are transgenically driven by the same *cryaa* promoter with single genomic insertion [33], we predict that the aggregation of γD-I4F involves the formation of 1 to 1 complexes with the R49C. Experiments are underway to test this conclusion.

In addition to enhanced binding affinity, R49C, but not R116C, has been shown to undergo disulfide cross-linking at the mutation site [41] suggesting that the location of the cysteine is critical. In studies of binding to the client protein βB1-crystallin, formation of an αA-R49C dimer that involved a mixed α/β-crystallin intermediate was observed [41]. This dimer was strictly dependent on the labeling of bB1 at a single cysteine by a disulfide-linked fluorescent probe, suggesting that the formation of chaperone/client protein complex is an intermediate step [41]. Because human γ-crystallins contain multiple cysteines, we predict that the formation of the complex with R49C is stabilized by disulfide cross-links. Previous studies have shown that the R14C mutation in human γD-crystallin increases the propensity of intermolecular disulfide bond formation [56, 57]. Importantly, in the homozygote knock-in R49C mouse line, the mutant αA-crystallin was reported to be disulfide cross-linked although the construct contained a second cysteine, which could lead to different types of covalent dimers. Consistent with our conclusion that disulfide cross-linking is the primary factor in the deleterious effect of the R49C mutation, the αB-crystallin R120G mutant did not promote the aggregation of γD-I4F.

Finally, a critical parameter typically missing *in vitro* is molecular crowding, which in lens fiber cells, is predicted to substantially alter the thermodynamics of molecular interactions. Not only does the excluded volume effect stabilize the folded states of client proteins but it is also predicted to affect the subunit exchange of the α-crystallins [58, 59]. Equilibrium dissociation of these proteins is a mechanism of chaperone activation [23]. In this context, it is notable that the two mutations have a profoundly different effect on the oligomeric structure [11]. Our finding of distinctive effects of the two mutants emphasizes the necessity of challenging mechanistic conclusions obtained *in vitro* in a model organism and highlights the role of cellular context in shaping chaperone interactions with native client proteins.

## Acknowledgments

We thank Dr. Derek Claxton for critical reading of the manuscript and Abigail Poff for technical assistance.

## Author contributions

S.-Y. Wu and H. Mchaourab designed research; S.-Y. Wu and P. Zou performed research; S.-Y. Wu analyzed data; S.-Y. Wu and H. Mchaourab wrote the paper; S.-Y. Wu, P. Zou and S. Mishra contributed new reagents.

## Footnote

This work was supported by NIH grants, R01 EY12018 and P30 EY008126.

